# HeartBioPortal: an internet-of-omics for human cardiovascular disease data

**DOI:** 10.1101/487744

**Authors:** Bohdan B. Khomtchouk, Kasra A. Vand, William C. Koehler, Diem-Trang Tran, Kai Middlebrook, Shyam Sudhakaran, Or Gozani, Themistocles L. Assimes

**Affiliations:** Department of Biology, Stanford University, Stanford, CA, USA 94305; Department of Medicine, Stanford University School of Medicine, Stanford, CA, USA 94305; VA Palo Alto Health Care System, Palo Alto, CA, USA 94304; Quiltomics, Palo Alto, CA, USA 94306; School of Computing, University of Utah, Salt Lake City, UT, USA 84112

**Keywords:** Computational Biology, Genetics, Gene Expression, Regulation

## Abstract

Cardiovascular disease (CVD) is the leading cause of death worldwide, responsible for over 17 million deaths annually, a rate which outpaces even that related to cancer. Despite these sobering statistics, the state-of-the-art in computational infrastructure for the study of contemporary datasets related to CVD lags substantially behind that widely available in oncology, where improved data science and visualization methods have delivered publicly available comprehensive cancer genomics resources like Memorial Sloan Kettering Cancer Center’s cBioPortal^1,2^ and the National Cancer Institute’s Genomic Data Commons (GDC) Portal^3,4^. In our view, such portals do an outstanding job of transforming data from The Cancer Genome Atlas (TCGA) into logical data visualizations that provide additional biological insight. Developing a similar user-friendly computational platform for CVD could significantly lower the barriers of discovery by providing researchers with rapid, intuitive, and high-quality visual access to molecular profiles and clinical attributes from existing CVD projects.

To our knowledge, no open-access computational resource for interactive visual exploration of multidimensional -omics datasets focused on CVD exists that rivals the utility, simplicity, and power of cBioPortal or the GDC Portal. While a large amount of data from various CVD studies has been deposited in dbGaP, these data are only accessible to researchers after a time-consuming manual review of databases to identify relevant datasets and a somewhat onerous administrative data request process. Within the field of CVD research, the American Heart Association’s Precision Medicine Platform (PMP) has significantly facilitated access to such data by streamlining the search, request, and transfer of controlled-access data and harmonizing datasets for cloud-based analyses^5^. Notwithstanding these important contributions of the PMP, we note that the Platform’s output currently mimics the content of TCGA (i.e., raw and harmonized data with no preprocessing or visualization of its internals). While the strength of the PMP centers on providing curated CVD datasets, programming tools/tutorials, and forums for collaboration, a need that remains unmet is an informatics infrastructure that can quickly distill CVD-related datasets into easy-to-interpret, insightful figures and charts in a manner analogous to the cBioPortal and GDC Portal, but whose content is updated on a regular basis.

Here we present HeartBioPortal, a publicly available web application that takes the first important steps towards fulfilling this need by integrating existing CVD-related omics datasets in real-time across the biological dataverse to provide intuitive visualization and analyses in addition to data downloads. The complementary focus of the HeartBioPortal platform is to provide community support for issuing gene/disease/variant-specific requests and visualizing the search results in a multi-omics context. By establishing an output like that of cBioPortal or the GDC Portal – containing preprocessed/analyzed data plus interactive visualizations accessible to the broader CVD and stroke research communities – we hope that HeartBioPortal significantly lowers the barriers between complex CVD datasets and researchers who require rapid, intuitive, and high-quality data visualization of molecular profiles and clinical attributes embedded within these datasets.

Currently, HeartBioPortal features gene expression, genetic association, and ancestry allele frequency information for 6005 variants in 22827 human genes across 12 broadly defined cardiovascular diseases spanning at least 243 research studies. A workflow architecture diagram is depicted (Figure). From a biological database perspective, HeartBioPortal currently syncs CVD-relevant data in real-time across at least the following publicly available resources: ClinVar, NHGRI-EBI GWAS catalog, OMIM, dbGaP, GTEX, CREEDS, HapMap, 1000Genomes, ExAC, TOPMed, gnomAD, Ensembl, and GEO. We coin the phrase “internet-of-omics” (an omics-centric hybrid of the internet-of-things (IoT) and internet-of-data fields) to describe the methods we’ve deployed to achieve such heterogeneous database interoperability. We believe the phrase appropriately emphasizes the challenges of extracting relevant biological information, in this case cardiovascular disease omics datasets, across various biomedical resource facilities (e.g., PubMed publications, metadata entries of data files stored in biological databases, etc.), where we have abstracted the concept of “things” from hardware (devices) to software (databases). The overarching goal of our “internet-of-omics” algorithms is to establish massively integrated e-connectivity of omics datasets from multiple biological data silos linked to each other by communication networks based on context. Harmonized tidy data downloads are provided directly through HeartBioPortal’s application programming interface (API) to support the data needs of academic clinical/laboratory CVD researchers while facilitating community transparency/cross-validation of the original data sources.

Future directions in the development of HeartBioPortal include substantially enriching the current offering of studies focused on gene expression by refining the phenotype definitions and integrating alternative splicing information (e.g., differential transcript usage and isoform-level expression). We also plan to expand HeartBioPortal’s genetic association content (currently restricted to the NHGRI-EBI GWAS Catalog) to include diverse GWAS consortium efforts as incorporated in the Broad Institute’s Cardiovascular Disease and Cerebrovascular Disease Knowledge Portals. The end goal is to facilitate exploratory data analysis of integrative multi-omics CVD data spanning a diverse trove of expression, association, and population genetic information.

A short demo of HeartBioPortal is provided here: https://www.youtube.com/watch?v=VLN3IRLGbHw&feature=youtu.be

**Figure.**
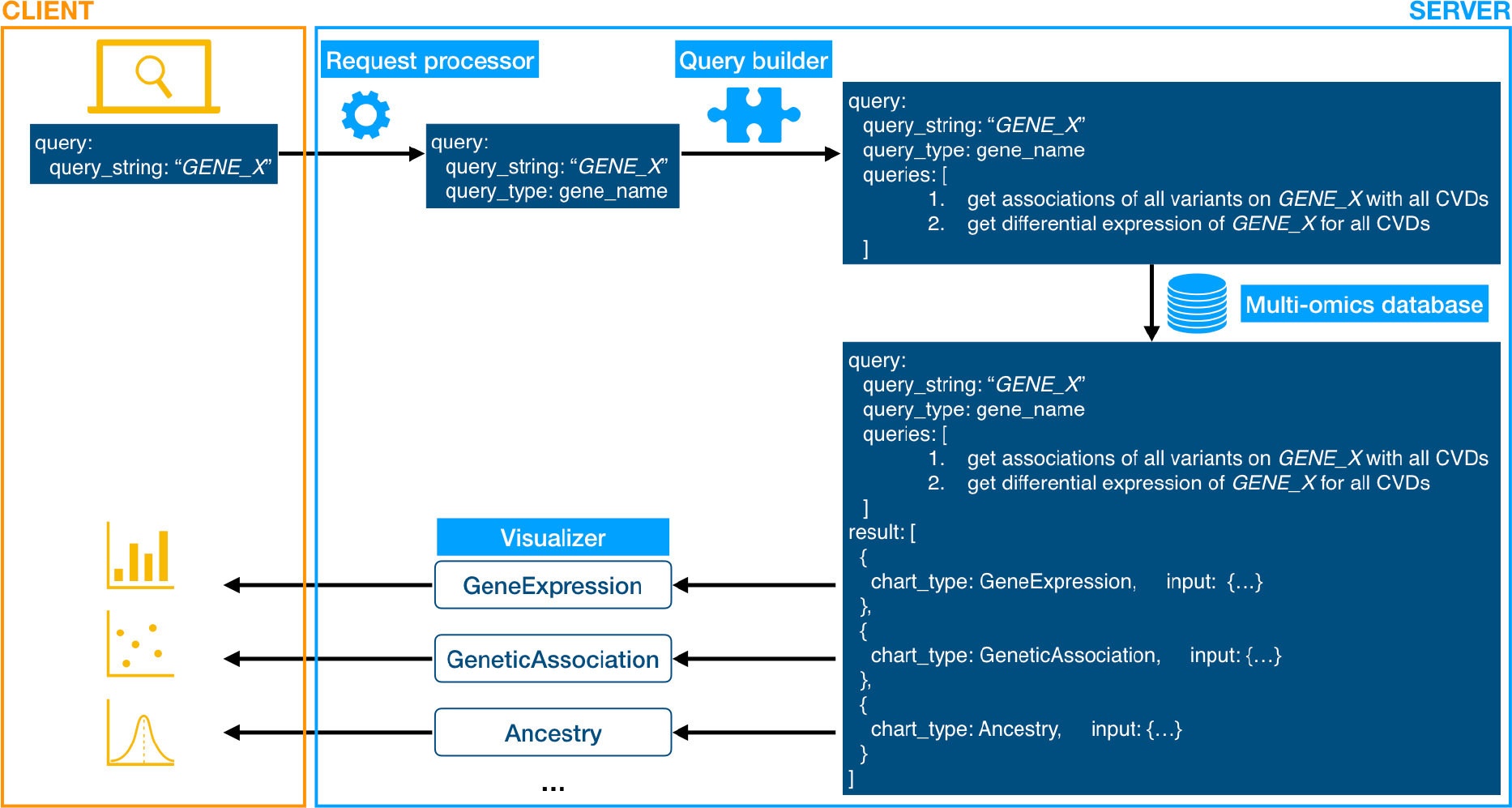
HeartBioPortal workflow architecture diagram. HeartBioPortal features a simple user interface that accepts queries for gene (current), disease/variant (future). User’s input is processed through multiple stages on the server into queries to be run on an integrated database, and eventually into preprocessed data that can be visualized in a meaningful way. Visualization components are highly extensible as the database grows to include larger and more diverse sources of omics data.

## Acknowledgements

BBK acknowledges and thanks the American Heart Association (AHA) for financial support through the AHA Postdoctoral Fellowship program. BBK thanks Komal Vadnagara and Evan Kaeding for useful discussions.

## References

1. E. Cerami, J. Gao, U. Dogrusoz, B.E. Gross, S.O. Sumer, B.A. Aksoy, A. Jacobsen, C.J. Byrne, M.L. Heuer, E. Larsson, Y. Antipin, B. Reva, A.P. Goldberg, C. Sander, N. Schultz: “The cBio Cancer Genomics Portal: An Open Platform for Exploring Multidimensional Cancer Genomics Data.” Cancer Discovery. 2012; 2(5): 401–404.

2. J. Gao, B.A. Aksoy, U. Dogrusoz, G. Dresdner, B. Gross, S.O. Sumer, Y. Sun, A. Jacobsen, R. Sinha, E. Larsson, E. Cerami, C. Sander, N. Schultz: “Integrative Analysis of Complex Cancer Genomics and Clinical Profiles Using the cBioPortal.” Science Signaling. 2013; 6(269): pl1.

3. R.L. Grossman, A.P. Heath, V. Ferretti, H.E. Varmus, D.R. Lowy, W.A. Kibbe, L.M. Staudt: “Towards a Shared Vision for Cancer Genomic Data.” The New England Journal of Medicine. 2016; 375: 1109–1112.

4. M.A. Jensen, V. Ferretti, R.L. Grossman, L.M. Staudt: “The NCI Genomics Data Commons as an engine for precision medicine.” Blood. 2017; 130: 453–459.

5. T.A. Kass-Hout, L.M. Stevens, J.L. Hall: “American Heart Association Precision Medicine Platform.” Circulation. 2018; 137: 647–649.

